# Between- and Within-subject Variability in Actigraphy Data: A Case Study on Sleep Patterns in Schizophrenia

**DOI:** 10.64898/2026.02.06.704499

**Authors:** Jinyuan Liu, Hanchang Cai, Zhichao Zhang, Jiawei Wang, Ellen Lee, Xinlian Zhang

## Abstract

Variability in longitudinal health outcomes provides critical insight into disease dynamics, yet most statistical analyses continue to prioritize mean trajectories. This limitation is particularly consequential in psychiatric and mobile health (mHealth) research, where intensive repeated measurements increasingly reveal pronounced heterogeneity and temporal instability that cannot be captured by average responses alone. Standard mixed-effects models typically impose homoscedastic variance assumptions and relegate variability to a nuisance component, despite mounting evidence that both variability information, at both between- and within-subject level, carry independent clinical significance. Here, we describe an improved mixed-effects location–scale model for longitudinal outcomes that simultaneously characterizes mean structure, between-subject heterogeneity, and within-subject fluctuations. The proposed framework explicitly links variance components to both time-invariant and time-varying covariates, extending classical mixed-effects models to accommodate heteroscedastic and dynamically evolving variability. By jointly modeling location and scale parameters, the method enables direct inference on variability (or stability) of the data alongside mean trends. We evaluated the approach using an actigraphy-based mHealth case study of sleep patterns in schizophrenia. Our results demonstrate that explicitly modeling variability uncovers clinically relevant structure obscured by mean-based analyses, underscoring the importance of variability-aware modeling for modern longitudinal and mHealth research.

## 1 Introduction

Variability in health and disease outcome trajectories, particularly in longitudinal psychiatric studies, represents a critical yet underexamined dimension of complex disorders such as schizophrenia. Traditional analyses have largely emphasized mean responses [1,2]. While group-level mean trajectories are intuitive and clinically interpretable, they often mask substantial inter- and intra-individual heterogeneity that characterizes psychiatric phenotypes.

A growing body of evidence suggests that such variability is by itself clinically meaningful. In schizophrenia, for example, mood swings, erratic sleep patterns, or inconsistent medication adherence have been linked to symptom exacerbation, relapse risk, cognitive impairment, and functional outcomes [3–5]. Individuals with similar mean sleep duration can differ markedly in stability, with implications for prognosis and intervention [6–8].

Standard statistical approaches in psychiatric research, including linear mixed-effects models (LME), typically focus on modeling mean structures while assuming homoscedastic variance across individuals and time [9]. Such over-simplified assumptions can obscure meaningful relationships between variability and key covariates. More flexible frameworks, such as mixed-effects location-scale models (LSM) that we focus on elaborating and generalizing here in this paper, directly address this limitation by jointly modeling the mean and variance components of longitudinal outcomes. These models allow both between-subject heterogeneity and within-subject fluctuations to be linked to clinically meaningful predictors, thereby providing a more nuanced characterization of psychiatric processes.

Recent advances in digital phenotyping further facilitate the proper modeling of variability. Technologies such as actigraphy and ecological momentary assessment (EMA) enable high-frequency, longitudinal measurement of behavior and physiology, yielding rich data on temporal instability at an unprecedented resolution [10]. While these data offer new opportunities to study dynamic processes, they also demand statistical methods that move beyond mean-based inference to fully exploit the information contained in patterns of variability.

The primary goal of this study is to enhance our understanding of variability in longitudinal psychiatric research by modeling both between- and within-subject variability in actigraphy data. Specifically, we aim to: 1) Discern the distinct sources of variability (between-subject heterogeneity, within-subject fluctuations, and measurement error) and evaluate their relative contributions to clinically relevant outcomes; and 2) Identify covariates that influence each variability component, including both time-invariant (e.g., diagnostic group) and time-varying ones (e.g., daily step counts).

## 2 Motivation

Longitudinal studies collect repeated measurements from the same individuals over time, which enable the assessment of changes over time and differences across subjects. A defining feature of such data is the within-subject correlation over time, which may be due to the shared underlying biological or behavioral processes [9].

Historically, the primary focus of longitudinal data analysis has been the mean trajectories or response profiles over time. In contrast, variability is often treated as a nuisance. But as already elaborated, variability parameters themselves bear potential to characterize stability, regulation, and heterogeneity of outcomes that are of great clinical relevance. In longitudinal settings, variability arises from three primary sources: between-subject heterogeneity, within-subject variability, and measurement error. We will clarify each of them in detail below.

Between-subject heterogeneity refers to the systematic differences in underlying response profiles across individuals. For example, in sleep studies, schizophrenia (SZ) patients may consistently exhibit longer sleep duration than healthy controls (HC), leading to separation in subject-specific trajectories. Such *inter*-individual heterogeneity motivates using subject-level random effects in mixed-effects models. In contrast, within-subject variability captures temporal fluctuations around an individual’s latent trajectory. Outcomes such as blood pressure, cortisol, and heart rate commonly exhibit such intra-individual dynamics, which are often biologically meaningful [6,7]. Measurement error, at last, represents random deviations of observed values from the true underlying process.

In the following discussions, we focus on the first two sources and assume measurement error is attenuated through improved data collection tools (e.g., actigraphy). The relative magnitude of within-subject and between-subject variability is critical for determining the degree of correlation among repeated measures. Under a compound symmetry assumption on the within-subject correlation, this relationship is summarized by the intra-class correlation coefficient (ICC),

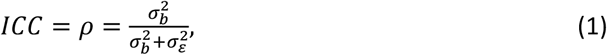

where 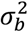 represents the between-subject variance, and 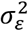 represents the within-subject variance. The definition in (1) reflects how the relative magnitude of two variance components contributes to the magnitude of temporal correlation.

### 2.1 One example: the mHealth Data

The mHealth data usually collect data at a much higher frequency or within a much intensive short-term period than traditional longitudinal studies [11–13]. This increased temporal resolution is enabled by wearable devices and smartphone-based assessments, allowing fine-grained measurement of time-varying behavioral and physiological processes [14,15].

Rich mHealth data sources include Ecological Momentary Assessment (EMA), which captures real-time psychological, social, and behavioral states [16–18], as well as large-scale electronic health record initiatives such as the UK Biobank [19,20] and *All of Us* [21,22], which incorporate daily activity metrics. These data offer unprecedented opportunities to study variability and its determinants at the individual level.

As a motivating example, consider an mHealth study in which SZ patients and HC participants wore wrist-based actigraphy devices continuously for 14 days to assess sleep and activity. Preliminary visualizations revealed that SZ patients demonstrated higher variability for the outcome of *sleep time*. It is of further interest to further distinguish the sources of variability and identify factors that affect each variability component.

The mHealth research increasingly treats variability itself as a primary outcome. Within-subject fluctuations may reflect clinically relevant dynamics driven by time-varying factors such as medication use or daily stressors [10]. Moreover, a growing body of evidence links elevated variability to adverse health outcomes: blood pressure variability predicts stroke risk [6,23,24]; mood instability is associated with stress [17], substance use [25], and depressive symptoms [6]; and cortisol intra-individual variability has been linked to major depressive disorder and autism [7,26].

However, statistical methods that explicitly model variability remain underdeveloped in psychiatric mHealth research. In this work, we position variability as a central inferential target. By explicitly decomposing and modeling sources of variability, we aim to evaluate how both time-invariant and time-varying factors shape variability components and how these effects evolve over time.

## 3 Methods

The mainstream linear mixed-effects models (LME) in the longitudinal setting can explicitly distinguish *between-subject* (BS) and *within-subject* (WS) variabilities. Consider a classical longitudinal study design with *N* subjects and *K* measurements, the LME with a random intercept models an outcome of interest *y*_*ik*_ for subject *i* (*i* = 1,2, …, *N*) at measurement *k* (*k* = 1,2, …, *K*) with

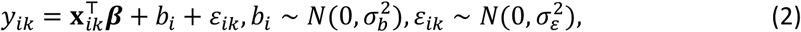

which designates 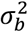 the between-subject variance and 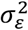 the within-subject variance, both assumed *homoscedastic* across groups or levels of covariates.

In mHealth studies, with more frequent measures per subject, the homoscedastic variance assumption may not hold, and interest oftentimes centers around the stability ot variability of outcomes. To this end, the mixed-effects location scale model (LSM) can be used to further model variance components:

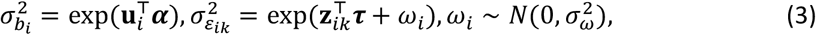

where exp(·) ensures correct range in modeling the variance. The changes in the two variance components are explained by time-invariant covariates **u**_*i*_ and time-varying covariates **z**_*ik*_. Now both 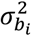 and 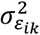 are allowed to differ across individual covariates, and further, 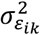 is allowed to vary across time. By parsing out the variance components, the covariate effects on variances can be described through parameters {**α, τ**}.

Formally, the LSM is a multi-level model specifying both the mean and variance components with

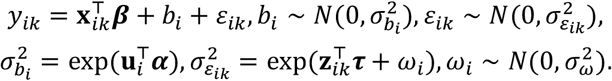

Unlike the mixed-effects model, it involves two random effects, with *b*_*i*_ the random *location effect* that influences an individual’s mean or location (*y*_*ik*_), and *ω*_*i*_ the random *scale effect* that impacts an individual’s variance or scale (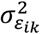).

Sometimes, the parameter **α** in between-subject variability is not of major interest; one can equivalently incorporate **u**_*i*_ into the random location, where the model simplifies to

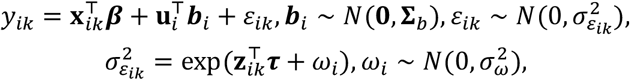

rendering parameters 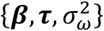 for estimation and inference. The estimation is through maximum likelihood estimation (MLE) while incorporating empirical Bayes methods for the random effects. The MixWILD in SAS and brms() function in R (R stan) are readily available.

## 4 Case Study

### 4.1 Dataset and Study Design

We illustrate the necessity and benefit of modeling variability components in our motivating mHealth study using actigraphy data. To investigate sleep patterns and physical activity in individuals diagnosed with schizophrenia (SZ) and healthy controls (HC). As summarized in **Supplemental Tab.1**, the total variability in daily sleep time among HCs spiked on the first day and subsequently decreased, whereas SZ participants consistently exhibited substantially higher variability, approximately twice that observed in HCs.

### 4.2 Statistical Modeling

To facilitate interpretation, we first focused on the first four days of data (all complete) then generalized to seven days as sensitivity analysis. The outcome of interest was daily sleep time (*y*_*ik*_), with diagnostic group (SZ vs. HC) as the main predictor (*x*_*i*_), and daily step count as a time-varying covariate (*z*_*ik*_).

The mean profiles indicated that HCs maintained relatively stable sleep times, while SZ participants started lower on Day 1, showed a spike on Day 2, and then decreased thereafter (see **Supplemental Fig.1**). Step counts followed a similar trend: low on Day 1, peaking on Day 2, which suggests a possible relationship between physical activity and sleep (**Supplemental Fig.2)**. A loess curve confirmed a non-linear association between daily step counts and sleep time (**Supplemental Fig.3**).

To disentangle between- and within-subject covariate effects of step counts, we applied person-mean centering. Specifically, each individual’s mean daily step count across days 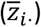 captured stable, between-person differences, while the daily deviation 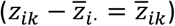 represented within-person fluctuations. After this partition, the non-linear trend diminished (see **Supplemental Fig.4**), suggesting that the earlier relationship was primarily driven by between-person differences.

#### 4.2.1 Linear Mixed Effects Model

We first fitted a standard linear mixed-effects (LME) model to evaluate mean sleep trajectories. The LME revealed a significant group-by-time interaction (omnibus *p* = 0.0271). No group difference was detected at baseline (*p* = 0.1433), but consistent with visualizations, the SZ group showed a significant increase in mean sleep time on Day 2 relative to HC (*p* = 0.0025).

Importantly, Supplemental Figure 1 and Supplemental Table 1 indicate that sleep duration in the SZ group displays substantially greater variability than in HC. This variability is not captured or explicitly modeled by the standard LME, which focuses primarily on conditional mean structure. Moreover, traditional longitudinal analyses do not disentangle the sources of variability, such as within-subject versus between-subject heterogeneity, thereby obscuring potentially meaningful differences in sleep stability across groups.

Further evidence of model misspecification was observed in diagnostic plots, which indicated violations of the homoscedasticity assumption (see **Supplemental Fig. 5**). Recall in LME from (2), random effects *b*_*i*_ should be independent, mean zero, with homoscedastic variance, and normally distributed. The residuals ε_*ik*_ should be independent, mean zero, constant variance, and follow a normal distribution. While random effects appeared normally distributed, the residuals exhibited clear heteroscedasticity across groups, with wider interquartile ranges and more dispersed residuals in the SZ group. The unusually large within- and between-subject standard errors further underscores the need for modeling frameworks that explicitly accommodate heteroscedasticity and variability structure, beyond what is permitted by a conventional LME.

#### 4.2.2 Location-Scale Model

To capture complex variability in sleep patterns, we extended the model to a mixed-effects location-scale model (LSM), which explicitly links mean and variance components to covariates. We illustrate the approach using both the 4-day dataset (complete data). We then try to approach the 7-day dataset with some missing data as sensitivity analysis.

We hypothesized that between-subject variance would depend on time-invariant covariates (e.g., diagnosis), while within-subject variance would be explained by time-varying effects.

##### Covariates

We considered the time-invariant covariates as follows:

1. Diagnostic Group (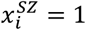 if SZ; 0 otherwise);
2. Personal Mean Step Count 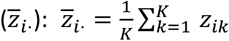.
3. Other key explanatory variables (**z**_*i*_), including stress (PSST), Physical Health, Positive and Negative Symptom Scores (SAPS, SANS), hba1c, age, and gender, were also considered in LSM.

Two time-varying covariates were considered:

1. Daily Step Count Deviation 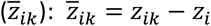
2. Day-by-Diagnosis Interaction 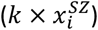, which allows for group-specific temporal effects.

##### Mean Response Model

We specify the mean sleep time (*y*_*ik*_) as a function of days, diagnostic group, and the day-by-diagnosis interaction, controlling for covariates (denote by **z**_*i*_).

For the 4-day sleep data, we view time as a discrete indicator and use dummy coding (**k**) considering the small number of days:

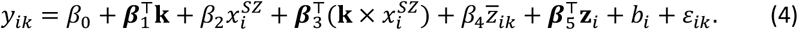

In the 7-day data, time was modeled flexibly using splines *s*(**k**):

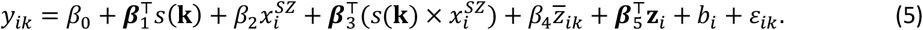

##### Variance Components Model

Between-subject variance was modeled as a function of diagnosis:

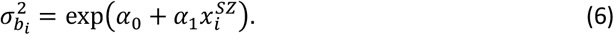

Lastly, we hypothesize that the within-subject sleep variability can be explained by day-by-diagnosis interaction and covariates. In particular, with the 4 day data, we posit:

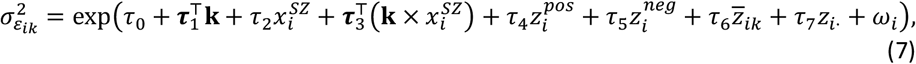

in the 7-day model, the categorical day terms were replaced with smooth functions:

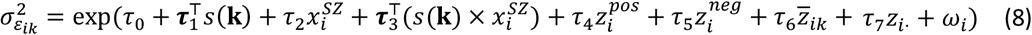

This hierarchical inclusion of covariates is helpful for investigators to disentangle the stable and dynamic influences on mean and variance components, yet to ease the computation, as in most software packages, one usually adopts the alternative form by incorporating 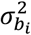 into the mean response model in (4) or (5) as a random slope:

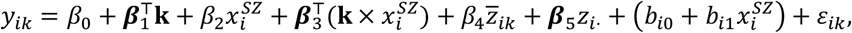

while keeping (7) or (8) unchanged, which include two sets of random effects with intercepts (*b*_*i*0_, *ω*_*i*_) and slope (*b*_*i*1_). Two sets of fixed effects, including ***β*** for location and **τ** for scale effects, will be estimated.

### 4.3 Results

#### 4.3.1 Model comparison: LME vs. LSM

The LSM model was implemented in brm() function in the bmrs package in R, paralleling the specification of lme4. **Supplemental Tab.2** compares the results between LME and LSM. As expected, fixed-effect estimates (***β***) exhibited mild shrinkage in LSM due to variance decomposition, while additional scale parameters **τ** and *σ*_*ω*_ provided explicit modeling of heterogeneity components. Hence, LSM identifies two conceptually distinct variance processes that are collapsed into a single composite variance under the LME.

#### 4.3.2 LSM Results with Covariates (4-Day Model)

We report the fixed effects (location and scale components) followed by the random effects, focusing on between- and within-subject variability in sleep time (see **Table 1**).

**Table 1:** Model results of LSM for 4-days, adding covariates, time as categorical.

| | Location ($\beta$) | Scale ($\tau$) |
| --- | --- | --- |
| Intercept | 332.65 | 4.42* |
| Day2 | 5.43 | -0.33 |
| Day3 | 9.32 | -1.81 |
| Day4 | 4.62 | -0.31 |
| SZ (vs. HC) | -68.78 | 0.27 |
| Personal-mean Step Count | -0.01 | 0.00 |
| Step Count Daily Deviation | -0.04 | 0.00 |
| Positive symptoms | -22.33 | 0.07* |
| Negative symptoms | -17.04 | 0.00 |
| Stress (psst) | -5.31 |  |
| Physical health (phycomp) | 1.52 |  |
| hbalc | -7.67 |  |
| Agevisit | 0.37 |  |
| Male (vs. Female) | -44.71 |  |
| Day2:SZ | 47.06* | 0.42 |
| Day3:SZ | 1.24 | 1.67* |
| Day4:SZ | 5.14 | -0.62 |
| $sd(b_{i0})$ | | 124.24* |
| $sd(b_{i1}) - SZ$ | | 351.64* |
| $cor(b_{i0}, b_{i1})$ | | 0.42 |
| $\sigma_{\omega}$ | | 0.53* |

\*indicates the effect falls in the 95% credible interval.

**Table 2:**
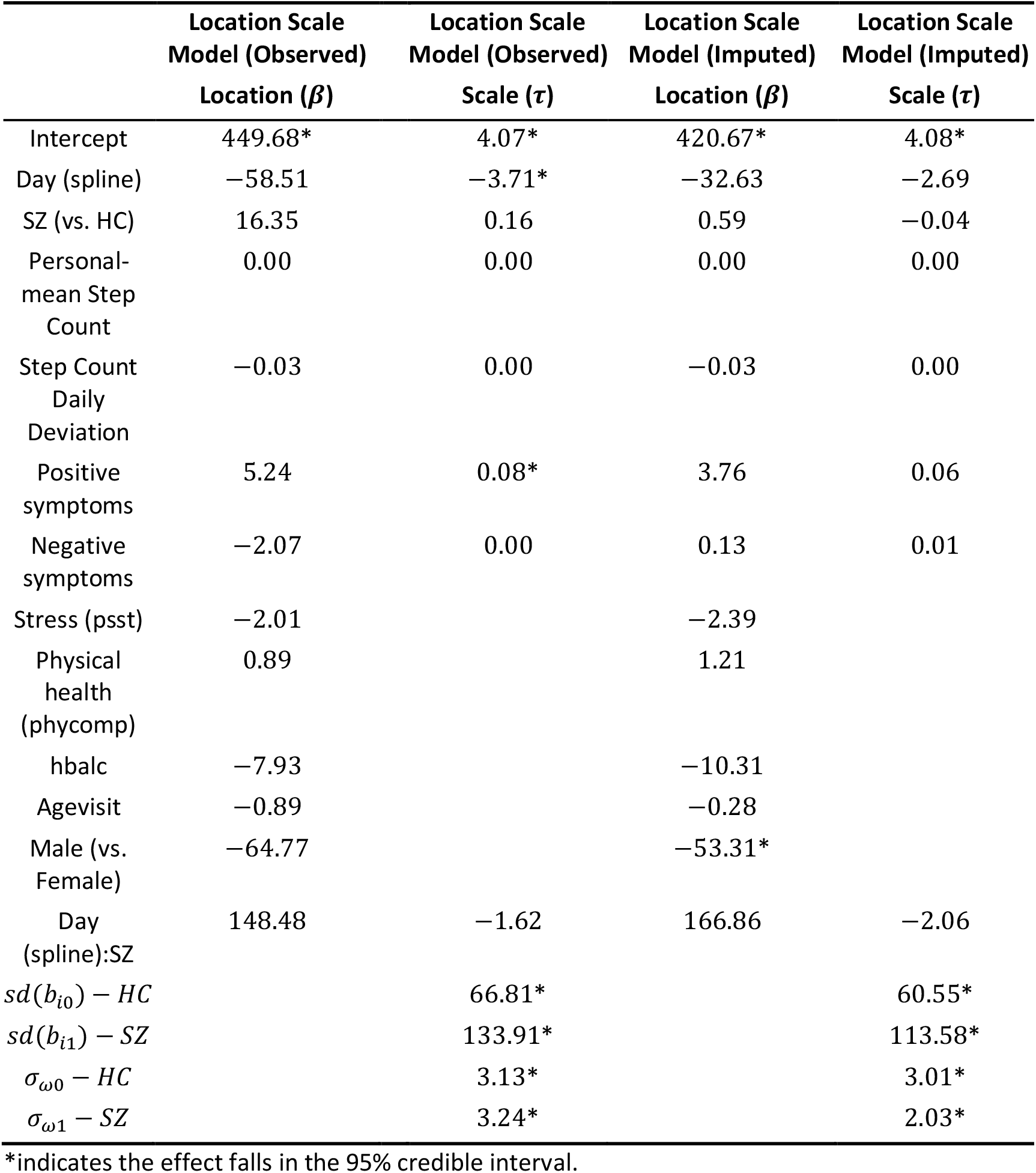
Model results of LSM for 7-days, adding covariates, time as spline (observed vs. imputed).

|  | Location Scale<br>Model (Observed) | Location Scale<br>Model (Observed) | Location Scale<br>Model (Imputed) | Location Scale<br>Model (Imputed) |
| --- | --- | --- | --- | --- |
| | Location ($\beta$) | Scale ($\tau$) | Location ($\beta$) | Scale ($\tau$) |
| Intercept | 449.68* | 4.07* | 420.67* | 4.08* |
| Day (spline) | −58.51 | −3.71* | −32.63 | −2.69 |
| SZ (vs. HC) | 16.35 | 0.16 | 0.59 | −0.04 |
| Personal-mean Step<br>Count | 0.00 | 0.00 | 0.00 | 0.00 |
| Step Count<br>Daily<br>Deviation | −0.03 | 0.00 | −0.03 | 0.00 |
| Positive<br>symptoms | 5.24 | 0.08* | 3.76 | 0.06 |
| Negative<br>symptoms | −2.07 | 0.00 | 0.13 | 0.01 |
| Stress (psst) | −2.01 |  | −2.39 |  |
| Physical<br>health<br>(phycomp) | 0.89 |  | 1.21 |  |
| hbalc | −7.93 |  | −10.31 |  |
| Agevisit | −0.89 |  | −0.28 |  |
| Male (vs.<br>Female) | −64.77 |  | −53.31* |  |
| Day<br>(spline):SZ | 148.48 | −1.62 | 166.86 | −2.06 |
| $sd(b_{i0}) - HC$ | | 66.81* | | 60.55* |
| $sd(b_{i1}) - SZ$ | | 133.91* | | 113.58* |
| $\sigma_{\omega 0} - HC$ | | 3.13* | | 3.01* |
| $\sigma_{\omega 1} - SZ$ | | 3.24* | | 2.03* |

\*indicates the effect falls in the 95% credible interval.

##### Fixed Effects Location (Mean) Fixed Effect

At baseline, SZ participants had lower mean sleep times than HCs (Estimate: -68.78), though not significantly. On day 2, both groups showed an increase in mean sleep duration, with SZ demonstrating a significant increase (Estimate: 47.06; 95% Credible Interval [3.39, 138.16]) compared to HC. Afterward, both groups continued to show moderate increases in mean sleep time; however, the between-group differences were not statistically credible. Step counts and stress measures showed negligible effects, while symptom scores (positive and negative) were weakly but negatively related to sleep time. Males tended to have shorter average sleep durations (Estimate: -44.71), but this effect was not significant.

##### Scale Fixed Effec

Within-subject variance differed by group and day, with SZ participants generally more variable than HCs. The group difference in variance reached statistical credibility on Day 3 (Estimate: 1.67, 95% Credible Interval [0.05, 5.03]), suggesting transient instability in sleep duration for SZ participants. Interestingly, higher positive symptom severity was also associated with increased within-person sleep variability (Estimate: 0.07, 95% Credible Interval [0.00, 0.15]). Step counts had no measurable effect on variance. Following prior recommendations [9,27,28], we interpret the random effects descriptively.

##### Location Random Effects

Considerable individual heterogeneity was found within both groups: the variance of subject-specific means was larger among SZ participants (Estimate: 351.64) than among HCs (Estimate: 124.24).

##### Scale Random Effects

The scale random effects capture subject-specific deviations in within-person sleep variability, thereby isolating individual differences in sleep stability that are unobservable under the LME. These effects indicate that participants differ not only in their average sleep duration but also in the magnitude of their within-subject fluctuations, even after accounting for observed covariates.

Overall, our model demonstrates SZ participants exhibited shorter and more variable sleep, with heterogeneity consistent with clinical observations. Stress and symptom measures exhibited expected but statistically nonsignificant associations with reduced mean sleep duration. Yet positive symptom severity was also associated with increased within-person sleep variability. Random-effects estimates underscored pronounced between-subject heterogeneity, especially among SZ participants. These conclusions are made possible through the implementation of modeling and disentangling the variability of data through out LSM.

#### 4.3.3 Sensitivity Analyses (7-Day Model: observed vs. imputed)

Due to attrition beyond seven days, multiple imputation was performed using the mice() package, and time was modeled as a spline.

Extending the model to 7 days yielded broadly consistent findings with the 4-day analysis. The general pattern of group differences and covariate effects also remained stable across both the complete-case and imputed datasets, with only minor changes in effect magnitude and precision.

As in the 4-day model, mean sleep duration did not differ significantly by diagnosis, and time effects were modest. Covariates such as step counts, stress, and symptom scores again had minimal impact on mean sleep time. Male participants consistently demonstrated shorter average sleep durations; this effect became statistically credible in the imputed model (−53 min vs. -65 min in the observed data).

Within-subject variability showed patterns nearly identical to the 4-day results: SZ participants remained more variable than HCs, with a mild decline in variance across days (negative coefficient in the spline). Positive symptoms remained the only covariate consistently associated with increased variability (Estimate: 0.08* observed; 0.06 imputed), replicating prior findings. Other predictors, including step counts and negative symptoms, again showed negligible effects. Between-subject heterogeneity remained higher in SZ (133.91 observed; 113.58 imputed) than in HC (66.81 observed; 60.55 imputed), which highlighted the robustness of group-specific variability.

Overall, the 7-day sensitivity analyses confirmed the primary findings of the 4-day model: schizophrenia was associated with greater intra-individual variability but comparable mean sleep duration to controls. The close correspondence between complete-case and imputed results strengthens confidence in the robustness of the LSM findings.

## 5 Discussion

In this paper, we discussed the flexible location-scale model (LSM) in exploring between- and within-subject variability in intensive longitudinal data, with a case study on sleep patterns in Schizophrenia.

The modeling framework employed by LSM offers several notable advantages. By conceptualizing the analysis to address both the mean response and variability components, this dual-focus framework allows for the disentanglement of mean effects and variability sources, offering richer insights into both group-level and individual-level patterns.

A key strength of this method lies in its ability to parse out exceptionally high volatility observed in mHealth or intensive longitudinal data. With datasets often comprising a short period or a large number of observations per individual, this approach is particularly suited for capturing nuanced patterns of variability that may otherwise be obscured in standard analyses. For instance, understanding within-subject variability can shed light on behavioral instability or treatment response fluctuations, while between-subject variability highlights heterogeneity across individuals. Additionally, this framework demonstrates greater flexibility than classical mixed effect model, which relies on homogeneous variance assumptions. By explicitly modeling variance components as functions of covariates, the proposed approach accommodates heteroscedasticity, providing a more realistic representation of the data and the processes underlying variability.

Despite its advantages, one key challenge of LSM is the greater number of parameters to estimate, which increases the risk of identifiability in smaller datasets. Linking variance components to covariates requires careful model specification and interpretation. Moreover, the reliance on parametric assumptions presents a potential limitation, as these assumptions, particularly the distribution of random effects, cannot always be validated in practice.

Computationally, the proposed framework may be difficult to scale to large datasets. Moreover, the Bayesian paradigm for hypothesis testing departs from classical frequentist approaches, which may complicate interpretation for practitioners more accustomed to traditional inferential frameworks.

As a future direction, a more robust and scalable alternative can be pursued within a semiparametric framework. For example, prior work has demonstrated the plausibility of using squared Euclidean distances to capture variability [29], which can recover coefficients analogous to those in the LSM while yielding more robust inference due to its semiparametric nature.

