## Supplementary Materials for "Between- and Within-subject Variability in Actigraphy Data: A Case Study on Sleep Patterns in Schizophrenia"

### Supplemental Fig.1: Mean sleep time trajectories across 4 days for healthy control (HC) and schizophrenia (SZ) groups, with error bars indicating standard deviations.

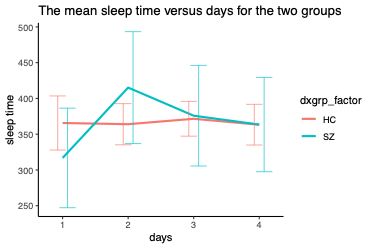

### Supplemental Fig.2: Mean daily step count trajectories across 4 days for healthy control (HC) and schizophrenia (SZ) groups, with error bars indicating standard deviations.

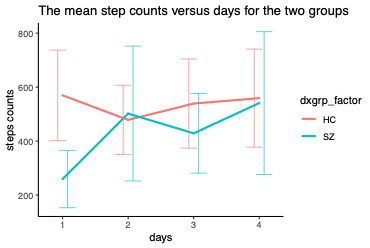

### Supplemental Fig.3: Loess smoothed curves showing the nonlinear association between daily sleep time and daily step counts for healthy control (HC) and schizophrenia (SZ) groups, with shaded regions representing 95% confidence intervals.

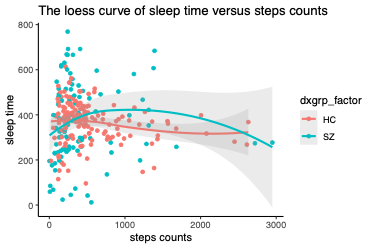

### Supplemental Fig.4: Association between daily sleep time and person-mean centered daily step counts (deviation from individual mean) for healthy control (HC) and schizophrenia (SZ) groups, with fitted linear trends and 95% confidence intervals.

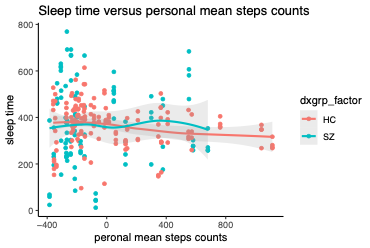

### Supplemental Fig.5: Diagnostic plots for linear mixed-effects model residuals showing (upper) standardized residuals by diagnostic group and (lower) standardized residuals versus fitted values for healthy control (HC) and schizophrenia (SZ) groups, demonstrating violation of the homoscedasticity assumption with greater residual variance in the schizophrenia group.

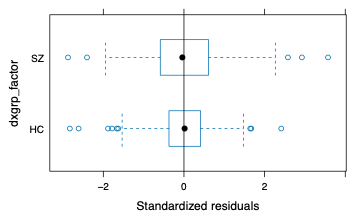

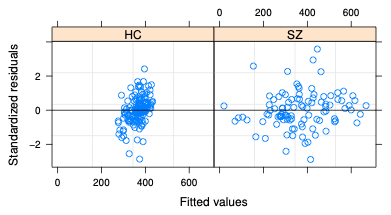

### Supplemental Tab.1: Total variabilities of *sleep time (min)* comparing two groups during the first four days.

| **Measurements** | $\mathbf{1}$ | $\mathbf{2}$ | $\mathbf{3}$ | $\mathbf{4}$ | **Mean** |
| --- | --- | --- | --- | --- | --- |
| HC | $130.16$ | $99.42$ | $83.42$ | $95.90$ | $102.2$ |
| SZ | $195.43$ | $221.25$ | $193.54$ | $181.73$ | $198.0$ |

**Supplemental Tab.2**: Model results comparison between linear mixed-effects model and location scale model.

|  | **Mixed Effect Model (**$\boldsymbol{\beta}$**)** | **Mixed Effect Location (**$\boldsymbol{\beta}$**)** | **Location Scale Model Scale (**$\boldsymbol{\tau}$**)** |
| --- | --- | --- | --- |
| Intercept | $367.77$* | $301.42$* | $4.50$* |
| Day2 | $-2.22$ | $4.05$ | $-0.39$ |
| Day3 | $5.71$ | $8.49$ | $-1.28$* |
| Day4 | $-2.43$ | $11.66$ | $-0.38$ |
| SZ (vs. HC) | $-54.92$ | $-39.19$ | $0.02$ |
| Personal-mean Step Count | $-0.01$ | $-0.01$ | $0.00$ |
| Step Count Daily Deviation | $-0.05$ | $-0.03$ | 0.00 |
| Day2:SZ | $102.02$* | $86.02$* | $0.49$ |
| Day3:SZ | $54.33$ | $33.78$ | $1.16$* |
| Day4:SZ | $50.74$ | $21.74$ | $-0.28$ |
| ${sd(b}_{i0})$ | $159$ |  | $67.07$* |
| ${sd(b}_{i1})-SZ$ | $142$ |  | $121.79$* |
| $cor(b_{i0}, b_{i1})$ | $-0.96$ |  | $0.20$ |
| $\sigma_{\omega}$ |  |  | $0.48$* |

* indicates passing 95% credible intervals for the model.
